# Evaluation of hippocampal *DLGAP2* overexpression on cognition, synaptic function, and dendritic spine structure in a translationally relevant AD mouse model

**DOI:** 10.1101/2025.05.13.653830

**Authors:** Andrew Ouellette, Kristen O’Connell, Catherine Kaczorowski

## Abstract

**INTRODUCTION:** Developing effective therapeutics for Alzheimer’s Disease (AD) requires a better understanding of the molecular drivers of the disease. Our previous work nominated *DLGAP2* as a modifier of age-related cognitive decline and risk for AD. We tested the hypothesis that overexpression of DLGAP2 in the hippocampus would protect against cognitive and synaptic deficits in a susceptible F1 5XFAD model.

**METHODS:** *DLGAP2* was overexpressed in the hippocampus of F1 hybrid 5XFAD and nontransgenic littermates using a viral approach. Cognitive function, electrophysiological properties, and dendritic spine morphology were assessed at 6 and 14 months of age.

**RESULTS:** *DLGAP2* overexpression impaired synaptic plasticity and exacerbated AD-related memory deficits but had minimal effect on spine structure or intrinsic neuronal properties.

**DISCUSSION:** We highlight the complex role of *DLGAP2* in AD pathology. Targeted interventions involving postsynaptic proteins must consider potential adverse effects on synaptic integrity and cognitive performance, particularly in the context of AD.

## 1. Background

Alzheimer’s disease (AD) is a progressive neurodegenerative disorder characterized by the accumulation of amyloid-beta plaques, tau tangles, and significant neuronal/synapse loss, ultimately leading to cognitive decline. While age is the most penetrant factor for the development of AD pathology, understanding the molecular underpinnings of the disease is critical for the development of targeted therapeutic strategies. Our previous cross-species analysis identified the postsynaptic density (PSD) protein *DLGAP2* (Discs-Large Associated Protein 2) as a potential mediator of AD-related cognitive outcomes and showed that ROSMAP AD patients with higher levels of DLGAP2 expression had improved late-life cognitive outcomes [1]. The *DLGAP2* protein is a key component in synaptic function and plasticity, organizing the postsynaptic density and mediating interactions between glutamate receptors and the actin cytoskeleton [2]. Additionally, *DLGAP2* has been implicated in neurological disorders such as autism spectrum disorder and schizophrenia [3–5]. However, its specific involvement in cognitive aging and AD remains poorly understood. Based on our previous work, we hypothesized that an increase in *DLGAP2* expression in memory-relevant brain regions would minimize or prevent cognitive deficits associated with AD neuropathology [1]. Therefore, in this study, we set out to test whether viral-mediated overexpression of *DLGAP2* in the hippocampus would improve cognitive outcomes in the 5XFAD.D2F1 mouse strain with the highest polygenic risk score for human AD as well as being associated with lower cognitive resilience to AD in mice [6, 7], and better understand the mechanisms in which *DLGAP2* may modify cognition and synaptic form/function.

## 2. Methods

### 2.1. Animals

The B6-5XFAD.D2F1 experimental mice and their nontransgenic littermates (B6.D2F1) were produced by trio mating female 5XFAD mice on a congenic C57BL/6J background (#034840, The Jackson Laboratory) with male DBA/2J (#000671, The Jackson Laboratory). Mice were group housed and maintained on a 12-hour light/dark cycle with ad libitum access to food and water. Mouse experiments occurred at The Jackson Laboratory in accordance with the NIH Guide for Care and Use of Laboratory Animals. All procedures and protocols were reviewed and approved by The Animal Care and Use Committee of The Jackson Laboratory (Animal use summary number: 16049). A total of 161 female mice were used for this study. The number of mice needed per group was derived from a power analysis performed in GPower 3.1 [8]; 9 mice per group are needed to detect a 10% difference in spontaneous alterations on the Y-maze task with ANOVA significance testing (alpha = 0.05, power = 0.85, 3 predictor variables and 8 groups). Based on our previous experience with surgical attrition and electrophysiology success rate this number was doubled.

### 2.2 Viral Constructs

Adeno-associated viral (AAV) vectors were produced by Vector Biolabs (Philadelphia, PA, USA). *Dlgap2* cDNA and eGFP controls were both packaged into AAV serotype 9 vectors and stored in a 1X PBS buffer containing 5% glycerol at a 1.1 x10^13^ titer. Protein expression in both vectors was driven by a CamKII promoter and had an HA tag fused to the C-terminus of the expressed protein. The resulting vectors were subjected to purification by two rounds of CsCl Density Gradient Centrifugation combined with ultracentrifugation to separate empty and full capsids.

### 2.3 Intrahippocampal injections

At 4mo of age mice were anesthetized under 4-5% inhalable isoflurane gas and positioned in a stereotaxic frame then maintained at 1-2% inhalable isoflurane gas. Two holes were drilled into the skull at the following coordinates: −1.9AP, +/− 1.4 ML, −1.6 DV. A microinjector attached to a 33-gauge Hamilton syringe and syringe pump were used to deliver 500 nL of virus per hemisphere at a rate of 200 nL/min. Microinjectors were left in place for 5 minutes following the end of injection to allow virus to diffuse adequately. Following injection, the cut was sutured using absorbable sutures. Post-surgery, topical bupivacaine was administered, and mice were monitored once daily for a minimum of three days. Viral spread was limited to the hippocampus/entorhinal cortex and did not spread to the cortex (Fig. S1).

### 2.4 Y-maze working memory task

All mice underwent the Y-maze spatial working memory task at 6mo of age, the aged cohort underwent a second trial at 14mo. Mice were allowed to freely explore a Y-maze apparatus with three equally sized arms for 8 minutes: arm length of 30 cm, arm width of 5-6 cm, wall height of 12-18 cm. Working memory was measured by % spontaneous alternations (Number of spontaneous alternations/Total arm Entries) between Y-maze arms. Recorded videos were analyzed using the ANY-maze behavioral tracking software (Stoelting Co., IL, United States). To limit the number of false positive arm entries we counted an arm entry when 99% of the mouse’s body (excluding tail) crossed into a new arm. We excluded data from 6 animals that did not make at least 6 total arm entries (the minimum number needed for at least 4 spontaneous alternations) from % spontaneous alternation data analysis. Distance traveled during testing (in meters) was also recorded as a metric of general locomotor activity.

### 2.5 Contextual fear conditioning

At 6 months of age all mice underwent Contextual Fear Conditioning (CFC) to assess the acquisition and recall of hippocampal-dependent memory [9]. On the first day of training, mice were placed in a training chamber and four foot-shocks (0.9 mA, 1 s) were delivered after a 150 second baseline period. Four post-shock intervals were defined as the 40 s following the end of each foot shock and the percentage of time spent freezing during each interval was determined using FreezeFrame software (Actimetrics Inc., IL, United States). The percentage of time spent freezing following the final shock was used as a measure of contextual fear acquisition across the panel. Twenty-four hours after training, mice were placed back into the training chamber and the percentage of time spent freezing throughout the entire 10-min test was measured as an index of contextual fear memory recall; no shocks were delivered during the testing session. Overall, 78 mice were sacrificed after testing at 6mo of age; 72 mice were aged until the 14mo timepoint. At 14mo of age, the aged cohort was placed into the conditioning chambers for 10 minutes with no foot shocks to determine that memory of their initial training regime had been extinguished. One week later these mice underwent the same training and test protocol as their initial 6mo timepoint. Outcomes from these tests were used as indicators of aged memory acquisition and recall.

### 2.6 Electrophysiology recordings

Mice were anesthetized with isoflurane before being sacrificed for electrophysiology recordings at 0-3 days after terminal CFC testing; LTP outcomes were tested to ensure they were not confounded by the delay between CFC and harvest (F(3, 89) = 1.17, p > 0.05). Brains were rapidly removed and placed in ice-cold cutting artificial CSF (aCSF) containing (in mM): 93 NMDG, 2.5 KCl, 1.4 NaH2PO4, 30 NaHCO3, 20 HEPES, 25 D-Glucose, 3 Myoinositol, 5 Sodium L-Ascorbate, 2 Thiourea, 6.1 NAC, 3 Sodium Pyruvate, 0.01 Taurine, 10 MgSO4.7H2O, 0.5 CaCl2.2H2O, pH adjust to 7.4 and osmolarity to 300mOsm. Acute transverse hippocampal slices (300µm) were cut with a vibratome (Leica, VT1000S) in ice cold cutting aCSF. Slices were incubated at 34℃ in carbogen bubbled cutting aCSF for 18 minutes. Slices were held at room temperature for 1-4 hours before recording in holding aCSF containing (in mM): 94 NaCl, 2.5 KCl, 1.4 NaH2PO4, 25 NaHCO3, 20 HEPES, 25 D-Glucose, 3 Myoinositol, 2 Thiourea, 5 Sodium L-Ascorbate, 6.1 NAC, 3 Sodium Pyruvate, 0.01 Taurine, 2 MgSO4.7H2O, 2 CaCl2.2H2O, pH adjust to 7.4 and osmolarity to 300mOsm. Whole-cell current clamp recordings were made in the CA1 pyramidal neurons at 34℃ under visual guidance of a video microscope (Olympus, Moment CMOS camera) using patch pipettes with resistance of 3-5 MΩ filled with potassium gluconate based internal solution (in mM): 126 K-Gluconate, 4 KCl, 10 HEPES, 4 MgATP, and 0.3 NaGTP, 10 Na-phosphocreatine, 0.2% biocytin, pH adjust to 7.4 and osmolarity to 290mOsm. Recordings were performed in an aCSF solution containing (in mM) at 34℃: 119 NaCl, 2.5 KCl, 2.5 CaCl.2H2O, 1 MgSO4, 1.25 NaH2PO4, 23 NaHCO3, 10 D-Glucose. A concentric bipolar stimulating electrode (FHC, Bowdoin, ME) connected to an A365 stimulus isolator (WPI, Sarasota, FL) was placed into the tissue along the Schaffer collateral 50μm-200μm away from the patched cell’s soma. A Multiclamp 700B amplifier, pClamp 11.2.1 software, and Digidata 1550B interface (Molecular Devices) were used to acquire data. Recordings were acquired at a 10 kHz sampling frequency and digitized at 20 kHz in current clamp mode. Neurons’ membrane potential was held at −67 mV. Series resistance and capacitance were monitored and compensated throughout recordings. Neurons with >40 MΩ series resistance were excluded. Changes in input resistance were measured by injecting 1s square step current into the soma ranging from −50pA to 50pA in 10pA intervals, input resistance was calculated in Easy Electrophysiology. Action potential threshold current was measured by injecting 2ms current step into the soma ranging from 200pA to 600pA; the minimum current needed to elicit an action potential was recorded as the threshold current. After-hyperpolarization (AHP) was initiated by a 25-action potential burst. mAHP was measured as the peak negative membrane potential relative to baseline and the sAHP was measured as the average negative membrane potential relative to baseline at 1–1.05 s after last brief current injection of protocol for triggering post-burst AHP [10].

Synaptic throughput was assessed with input/output curves by applying a standardized current ramp to the Shaffer Collateral starting at 25μA, proceeding to 50μA, then increasing up to 600μA at 50μA intervals every 20 seconds. Excitatory postsynaptic potentials (EPSPs, mV) were recorded in response to each stimulus. The current needed to elicit a 3-5 mV response was used for paired pulse ratio and LTP recordings. We then monitored EPSPs 5-min baseline period, evoking an EPSP every 20s. To induce synaptic plasticity, a theta-burst stimulation (TBS) protocol was used that consisted of theta-patterned synaptic activation (5 stimuli at 100 Hz) of proximal neuronal inputs repeated at 5 Hz for 3s. EPSPs were then evoked for 35 minutes every 20 seconds; the peak amplitude of each EPSP relative to baseline membrane potential was recorded. These peak amplitudes were aggregated into five-minute bins for analysis. To calculate the paired-pulse ratio (PPR), two stimulus pulses were delivered at 25-, 50-, 75, or 100-ms intervals and the PPR was measured by dividing each EPSP by the first.

### 2.7 Immunohistochemistry

Immediately after each electrophysiology recording was complete, the brain slice was placed into a 4% paraformaldehyde (PFA) solution for 24 hours and then transferred to 1xPBS containing 0.1% sodium azide. Slices were incubated with rabbit anti-HA antibody (Cell Signaling, #C29F4) and streptavidin-Alexa 633 (Thermo, #S21375) at 1:1000 dilution each for 24 hours at 4℃. After washing, slices were incubated in 1:500 goat anti-rabbit AlexaFluor 568 (Thermo, # A-11036) for 2 hours at room temperature. During the final wash prior to mounting, slices were incubated with 1:2000 DAPI for 10 minutes. Slices were mounted on Colorfrost Plus slides (Fisherbrand) with 0.36mm spacers and SlowFade Glass Antifade Mountant (Invitrogen). Slides were stored in the dark at 4℃.

### 2.8 Image Acquisition

Confocal microscopy was used to capture images of biocytin filled dendrites from patched CA1 pyramidal neurons. Imaging was performed by a blinded experimenter on a Stellaris 5 confocal microscope (Leica. Wetzlar, Germany), using a Plan Apo 63×/1.40NA oil-immersion objective. Three-dimensional z-stacks were obtained of secondary dendrites from filled neurons that met the following criteria: (1) within 80 µm working distance of microscope, (2) relatively parallel with the surface of the transverse section, (3) no overlap with other branches, (4) located between 50 and 150 µm from the soma. Up to 5 apical and basal dendrites were acquired per cell. Dendrite segment images were acquired in z-stacks with a step size of 0.13 µm, image size of 1024 × 512px, zoom of 4.8x, line averaging of 4x, and acquisition speed of 400hz; individual pixel size was 0.36 x 0.36um.

### 2.9 Dendritic Spine Reconstruction

Confocal z-stacks of dendrites were deconvolved using Huygens Deconvolution System (16.05, Scientific Volume Imaging, the Netherlands) using the following settings: deconvolution algorithm: GMLE; maximum iterations: 10; signal to noise ratio: 15; quality: 0.003. Deconvolved images were saved in .tif format. Deconvolved image stacks were imported into Neurolucida 360 (2.70.1, MBF Biosciences, Williston, Vermont) for dendritic spine analysis. For each image, the semi-automatic voxel scoop algorithm was used to trace the dendrite. All assigned points were examined to ensure they matched the dendrite diameter and position in X, Y, and Z planes, and were adjusted if necessary. Dendritic spine reconstruction was performed automatically using a voxel-clustering algorithm, with the following parameters: outer range = 5 µm, minimum height = 0.3 µm, detector sensitivity = 80%, minimum count = 8 voxels. The automatically identified spines were examined to ensure that all identified spines were real, and that all existing spines had been identified. If necessary, spines were added by increasing the detector sensitivity to 100% and manually identified. Merge and slice tools were used to correct errors made in the morphology and backbone points of each spine. Each dendritic spine was automatically classified as a thin spine, stubby spine, mushroom spine or filopodia spine based on constant parameters. Three-dimensional dendrite reconstructions were exported to Neurolucida Explorer (2.70.1, MBF Biosciences, Williston, Vermont) for branched structure analysis, which provides measurements of dendrite length; number of spines; spine length; number of thin, stubby, and mushroom spines, and filopodia; spine head diameter; and spine neck diameter, among other measurements. Spine density was calculated as the number of spines per 10µm of dendrite length. Values per mouse were calculated by averaging the values for all dendrites corresponding to that mouse, statistics were performed with mice as the independent sample. To assess apical and basal dendrites separately, values for either apical or basal dendrites were averaged separately per mouse, number of animals, cells and dendrites imaged are listed in Table 1.

**Table 1.** Sample n sizes are listed for the number of animals allocated to each group at the start of study, behavior, electrophysiology and dendritic spine measurements. CFC was longitudinally measured, therefore the 14mo cohort is included in the 6mo data. For Electrophysiology measurements n size is reported as Mouse n/Cells n. For Dendritic spine data, n sizes are reported as mouse n/cells n/dendrite n.

| Age (mo) | Genotype | AAV Status | Allocated Mice | CFC | Y-maze | Input/Output | LTP | PPR | AP Current Threshold | Input Resistance | AHP | Apical Dendrites | Basal Dendrites |
| --- | --- | --- | --- | --- | --- | --- | --- | --- | --- | --- | --- | --- | --- |
| 6 | Ntg | Control | 20 | 43 | 37 | 7/13 | 6/10 | 7/11 | 7/11 | 7/13 | 7/13 | 6/8/20 | 4/4/6 |
| 6 | Ntg | Dlgap2 | 19 | 41 | 36 | 6/12 | 4/7 | 6/10 | 6/13 | 6/12 | 6/12 | 6/6/12 | 3/3/8 |
| 6 | 5XFAD | Control | 20 | 35 | 32 | 7/12 | 5/7 | 7/12 | 7/12 | 7/12 | 7/12 | 5/6/25 | 5/6/15 |
| 6 | 5XFAD | Dlgap2 | 19 | 35 | 33 | 8/12 | 6/9 | 8/12 | 8/12 | 8/12 | 8/12 | 8/10/32 | 6/8/18 |
| 14 | Ntg | Control | 24 | 24 | 24 | 8/14 | 7/11 | 9/14 | 9/15 | 9/15 | 9/15 | 7/9/31 | 7/9/19 |
| 14 | Ntg | Dlgap2 | 26 | 24 | 24 | 11/20 | 11/19 | 11/20 | 11/20 | 11/20 | 11/20 | 10/15/57 | 10/12/31 |
| 14 | 5XFAD | Control | 16 | 13 | 13 | 9/16 | 9/16 | 8/13 | 9/16 | 9/16 | 9/16 | 6/9/30 | 6/8/14 |
| 14 | 5XFAD | Dlgap2 | 17 | 11 | 11 | 8/13 | 8/11 | 7/10 | 8/14 | 8/13 | 8/13 | 7/9/30 | 4/5/11 |

## 3. Results

### 3.1 Experimental design and AAV Validation

An AAV-9 containing *Dlgap2* cDNA fused to an HA epitope behind a CamKII promoter was used to drive overexpression of DLGAP2 in neurons of the hippocampus (Fig. 1A-B). To verify that AAV successfully overexpressed *DLGAP2* in mice, we quantified *DLGAP2* protein expression via western blot of injected hippocampal samples. *DLGAP2* expression was significantly elevated in animals injected with *Dlgap2* AAV compared to eGFP controls at 14mo of age in both Ntg (mean change = 1.3, t(7) = −2.30, p <0.05) and 5XFAD animals (mean change = 1.1, t(6) = −3.01, p < 0.05) (Fig. 1C, Left, Fig. S2), indicating that *DLGAP2* is robustly overexpressed within the hippocampus. IHC in a subset of mice confirmed that AAV expression was largely in dorsal CA regions, Dentate Gyrus (DG), and Subiculum, and more ventral CA3. Expression was also observed along CA3-CA1 tracts in the medial hippocampus (Fig. 1C Right), but pyramidal cells originating from this layer.

**Figure 1.**
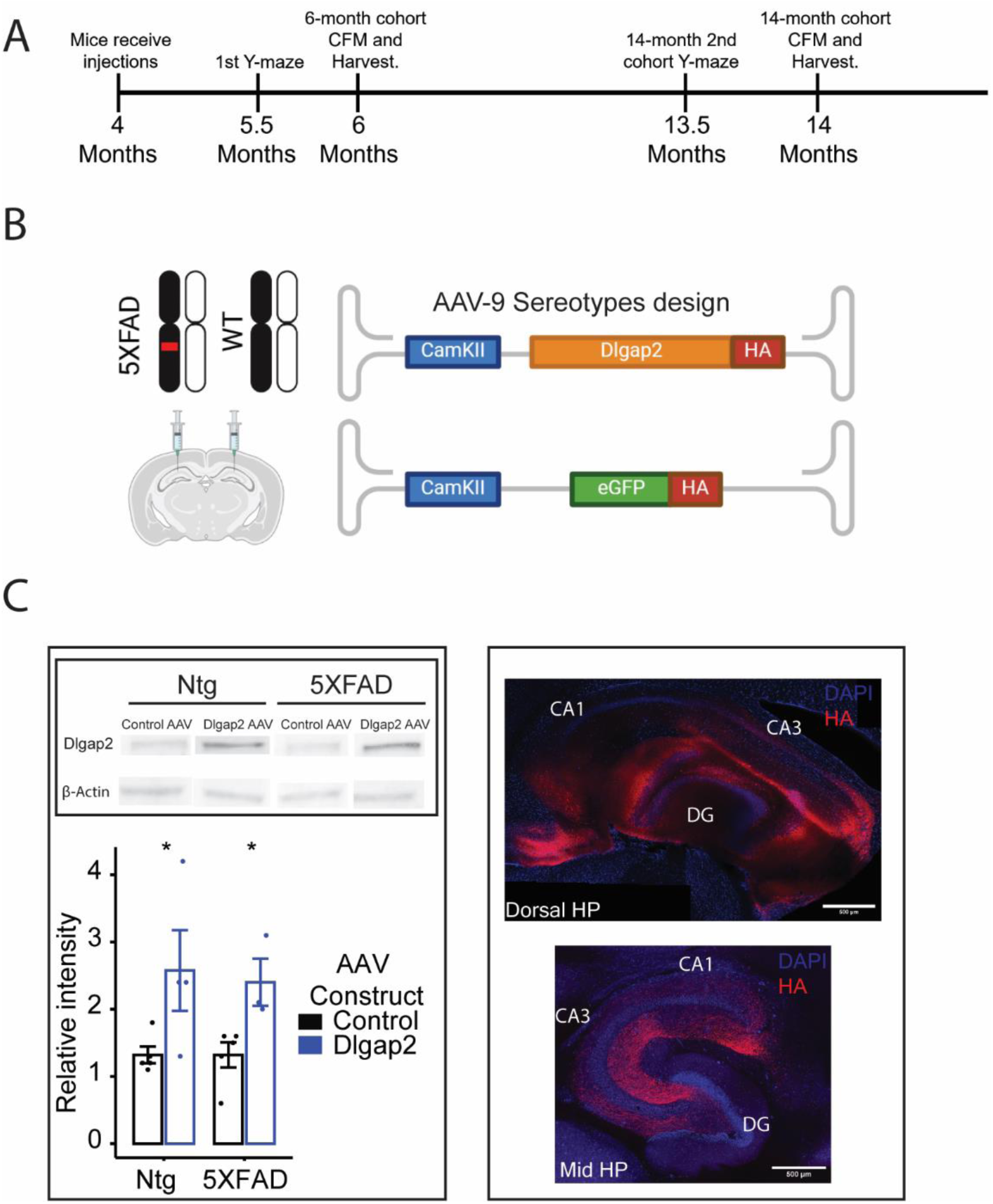
AAV Design and Study Overview. A) Timeline of experiments showing the age of mice in months starting at time of injection until terminal 6mo and 14mo timepoints. B) 5XFAD and WT animals were injected with an AAV-9 construct with either DLGPA2 or eGFP cDNA behind a CamKII promoter into hippocampus CA1. C) Left: Western blot results showing levels of *DLGAP2* protein expression in whole hippocampal sections from 14mo mice injected with either eGFP controls or *Dlgap2* cDNA. The intensity of *DLGAP2* bands is normalized to β-Actin loading controls, n = 3-5 per group. Right: Spatial visualization of exogenous *DLGAP2* in 14mo animals. In general, viral spread is mostly observed in dorsal CA regions, Dentate Gyrus and Subiculum, with highest expression seen in tracts projecting into more ventral CA1. *p<0.05.

### 3.2 DLGAP2 overexpression has no effect on spatial working memory

At 6 months of age, all mice were tested for spatial working memory using the Y-maze task; the aged cohort were retested again at 14 months. All groups performed above chance (50% SA), indicating non-random exploratory behavior. While we detected a statistically significant main effect of AAV construct on % spontaneous alternations (F(1, 27) = 5.98, p < 0.05), there were no significant post hoc comparisons after correcting for multiple comparisons (Fig. S3). These results, along with limited effect sizes, suggest that *DLGAP2* hippocampal overexpression does not significantly impact spatial working memory, regardless of age or AD status.

### 3.3 DLGAP2 overexpression is detrimental to AD-related contextual fear memory decline

To determine whether DLGAP2 overexpression can ameliorate AD-related cognitive deficits, we employed a novel longitudinal contextual fear conditioning paradigm at both 6 and 14mo of age. Age dependent memory acquisition and recall were assessed in all animals by measuring contextual fear acquisition (CFA) and contextual fear memory (CFM) respectively. We first established that initial training and testing in the 6mo and 14mo cohorts was comparable at their initial 6mo timepoint; three-way ANOVA indicates that both cohorts have comparable training/testing outcomes at 6mo (Fig. S4). Two-way repeated measures ANOVA detected a significant main effect of AD status on CFA where animals with the 5XFAD transgene tend to perform worse than Ntg counterparts especially by 14mo of age (F(1, 63) = 9.68, p < 0.01). We detected a significant interaction between AD status x AAV construct (F(1, 63) = 19.84, p < 0.001) where AD animals overexpressing *DLGAP2* have worse outcomes compared to control injections. We observed a similar pattern in CFM outcomes; two-way repeated measures ANOVA detected a significant decrease in *DLGAP2* overexpressing mice compared to control injections (F(1, 57) = 6.63, p < 0.05) and a decrease from 6mo to 14mo age points (F(1, 57) = 10.12, p < 0.01), alongside significant interactions between AD status x age (F(1, 57) = 23.78, p < 0.001) (Fig. S5), and AAV construct x age (F(1, 57) = 4.84, p < 0.05). At 6mo there was no significant effect of *DLGAP2* overexpression on CFA nor CFM outcomes (Fig. 2A-B). Interestingly, we observed an increase in CFM in 5XFAD control AAV animals compared to Ntg controls that was not observed in *DLGAP2* overexpressing animals

**Figure 2.**
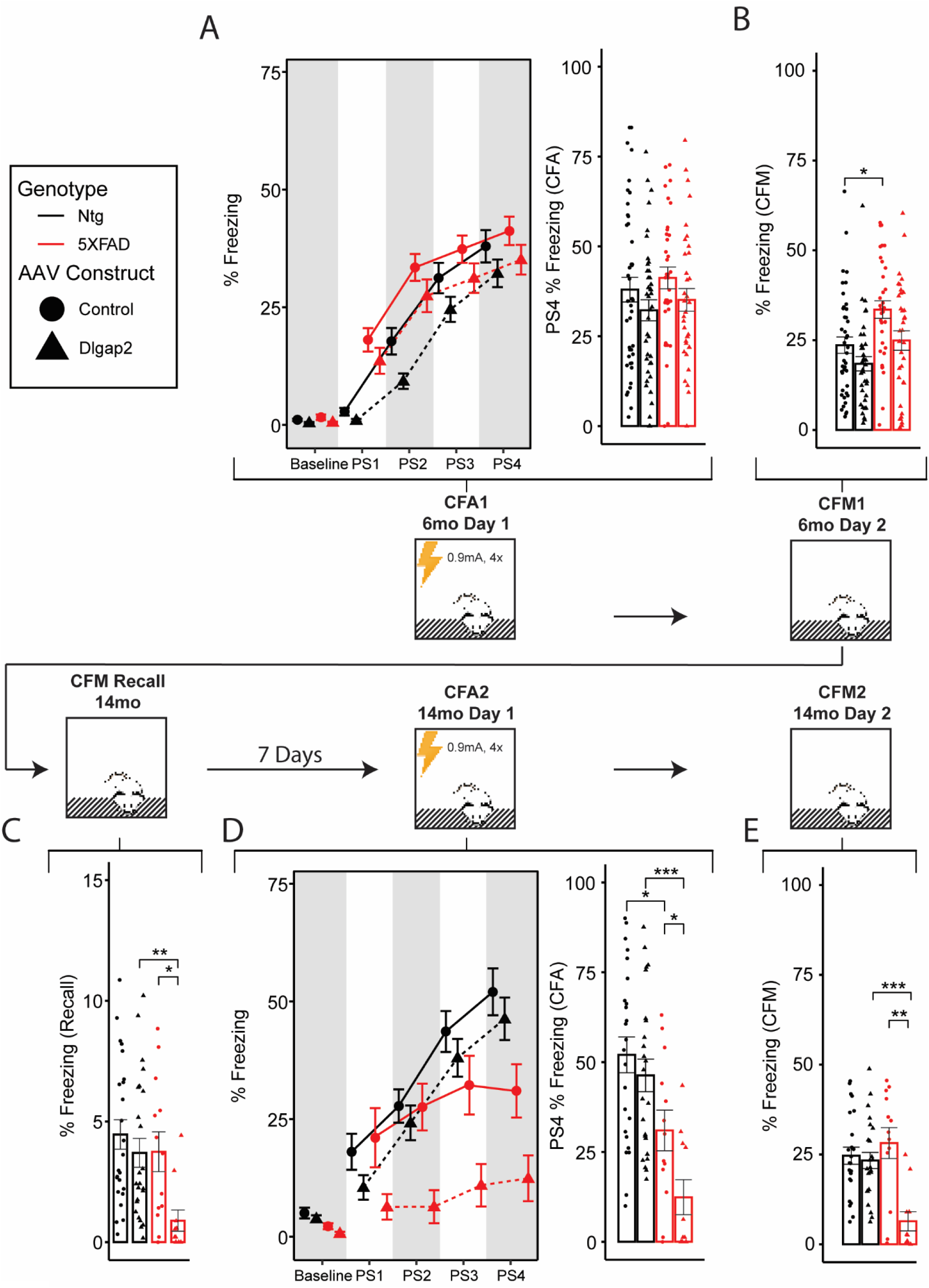
*DLGAP2* Overexpression impairs memory outcomes in aged 5XFAD animals. A) (Left) Memory acquisition curves measured by % time spent freezing during 150s baseline period before shocks were administered and each 40s post shock interval. (Right) 6mo contextual fear acquisition measured (CFA) as % time spent freezing during the 40s post shock 4 (PS4) interval. B) 6mo contextual fear memory (CFM) recall measured by % time freezing during the testing session with no shock administration 24hours after training. C) 7 days before 14mo retrain/testing, animals were placed into the chamber with no shocks and total % freezing was measured as an indicator of how well they remembered the initial 6mo training session. 5XFAD animals have a slight reduction in % freezing compared to controls. D) (Left) Memory acquisition curves during retraining of 14mo animals. (Right) CFA outcomes in 14mo animals during the retraining session; 5XFAD animals overexpressing *DLGAP2* have severe impairment in the ability to learn the task compared to controls. E) CFM outcomes in 14mo animals 24 hours after retraining. CFM was significantly impaired by *DLGAP2* overexpression in 14mo 5XFAD animals. Error bars are represented as standard error around the mean. p-values were derived from a t-test and corrected for multiple testing error using the Bonferroni method (*p<0.05, **p<0.01 ***p<0.001). n sizes are listed in Table 1.

To track individual changes in cognitive longevity, CFC was performed longitudinally in the 14mo cohort. Therefore, we tested if any experimental mice had retained memory of the initial 6mo testing phase and if this possible retention could affect retesting outcomes by measuring % freezing without foot shocks one week before terminal retraining/testing. Overall, animals had reduced memory recall compared to their 6mo testing session, which indicates that by 14mo of age animals do not retain the memory of their initial training. Additionally, we found that 5XFAD animals overexpressing *DLGAP2* had significantly lower memory recall than Ntg and control AAV counterparts, suggesting perhaps that *DLGAP2* overexpression impaired long term memory recall (Fig. 2C) (T(1,18)= 3.05, p adjusted < 0.05). However, the small effect size limits the interpretations that observed differences in CFA and CFM are confounded by prior memory retention.

By 14mo of age, 5XFAD animals displayed an expected decrease in CFA in mice compared to Ntg controls (T(1,21.9) = 2.48, p adjusted < 0.05) (Fig. 2D) [11]. However, we observed that *DLGAP2* overexpression was detrimental to CFA outcomes in 5XFAD animals. Slope acquisition curves show that aged 5XFAD animals overexpressing *DLGAP2* are unable to learn the context of their environment, and that their acquisition of the post-shock 4 interval was severely impaired (Fig. 2D). These data suggest that previously observed AD-related memory acquisition deficits are exacerbated with *DLGAP2* hippocampal overexpression. Additionally, overexpression of *DLGAP2* reduced CFM performance in 14mo 5XFAD animals compared to Ntg controls (Fig. 2E). In contrast to CFA, we did not observe a significant decrease in CFM in 14mo 5XFAD control AAV animals compared to Ntg controls, suggesting that *DLGAP2* overexpression accelerates AD-related impairment of CFM at 14mo of age. Since contextual fear conditioning outcomes can be confounded by changes in baseline locomotor activity similar to those we observed in the Y-maze [12, 13], we analyzed 150s of baseline freezing before shocks were administered on the training session across each experimental group. Two-way repeated measures ANOVA detected significant main effects of AD status (F(1, 63) = 8.36, p < 0.01), age (F(1, 63) = 29.3, p < 0.001) and an interaction between age x AD status (F(1, 63) = 14.85, p < 0.001); no significant main effect of *DLGAP2* overexpression was detected. Additionally, Post hoc analysis revealed that baseline freezing was reduced only in 14mo 5XFAD animals that had been injected with *DLGAP2* AAV compared to Ntg counterparts (Fig. S6B-F). Age also had a robust effect on baseline freezing where Ntg 6mo animals have significantly lower % freezing than 14mo in both control and *DLGAP2* overexpressing cohorts. This age-related change is not present in 5XFAD animals suggesting that AD pathology blunts the natural tendency for increased locomotor activity as animals mature. These results suggest that AD-related pathology may elevate locomotor activity and is exacerbated by *DLGAP2* overexpression but does not necessarily coincide with observed cognitive deficits.

### 3.4 DLGAP2 overexpression affects synaptic properties in pre symptomatic 5XFAD animals

DLGAP proteins play an important role in synapse form and function. To assess the relationship between synaptic function and structure during the progression of age-related AD pathology and *DLGAP2* overexpression, we characterized synaptic properties in patched CA1 pyramidal neurons and then imaged apical and basal dendritic spines. Specifically, we investigated whether overexpression of *DLGAP2* lead to alterations in synapse function by measuring synaptic throughput, paired pulse facilitation, and long-term potentiation (LTP) in CA1 pyramidal cells via stimulation of the Schaffer Collateral. We observed no significant effect of *DLGAP2* overexpression on synaptic throughput as measured by input/output curves (Fig. 3A). However, on average, brain slices from 14-month-old animals required higher stimulation to evoke comparable EPSPs to those from 6-month-old animals (F(1, 91) = 23.0, p < 0.01) (Fig. 3A). While 5XFAD animals overexpressing *DLGAP2* displayed comparable input/output curves at both 6 and 14 months, this effect was present in 5XFAD control injections and Ntg *DLGAP2* overexpressing animals. These results indicate that while *DLGAP2* overexpression does not directly modify synaptic throughput, there may be a synergistic effect of *DLGAP2* expression with AD pathology that minimizes age-related changes on the strength of synaptic signaling.

**Figure 3.**
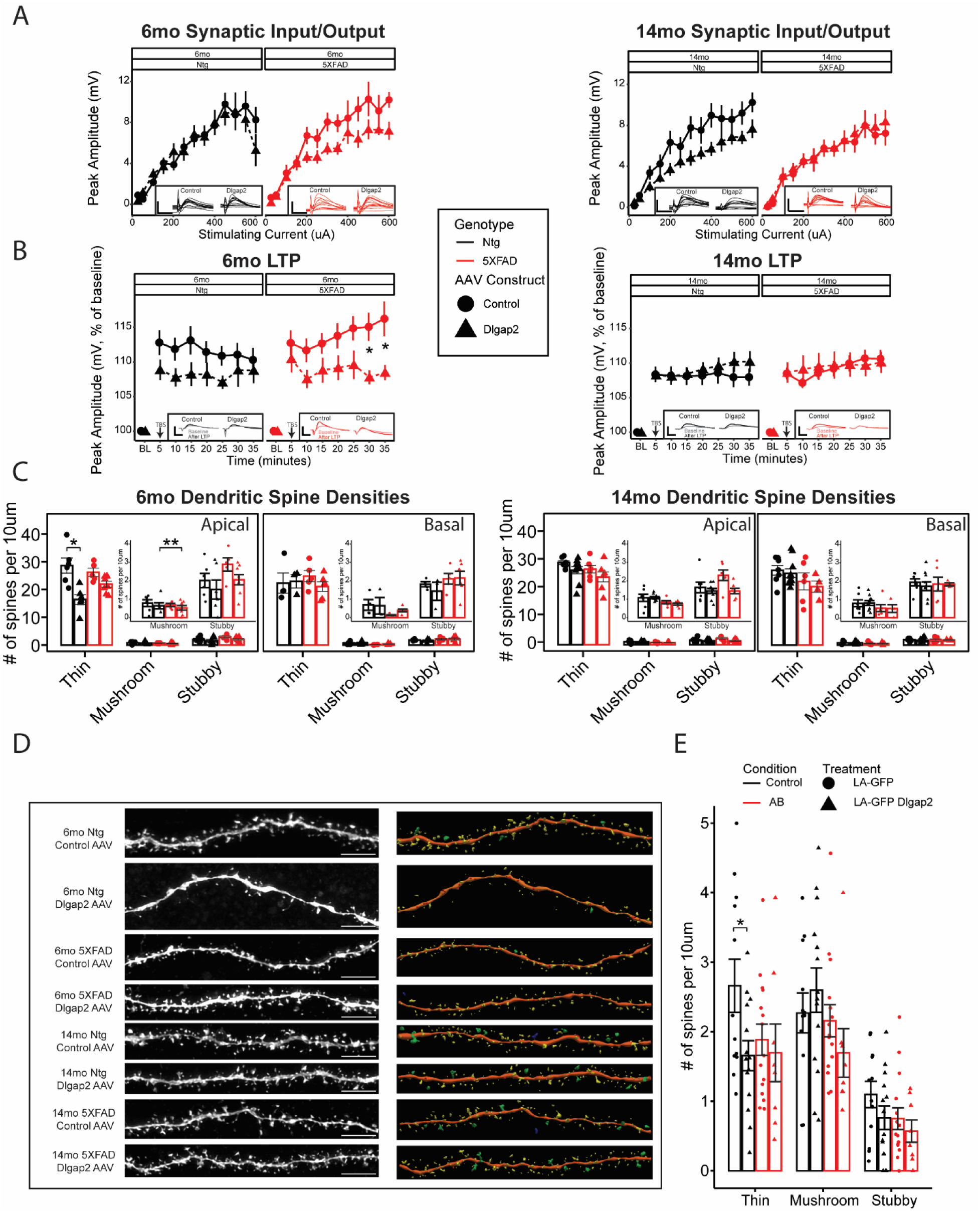
Synaptic functional and structural propeties exhibit *DLGAP2* overexpression, amyloid burden and age interactions. A) Input/Output curves, stimulating current injected into the schaffer collatoral is plotted against evoked epsp amplitude of patched CA1 pyramidal cells for 6mo (right) and 14mo (left) animals. Inset with representative traces (scale bar: 10ms,15mV), no main effect of *DLGAP2* overexpression was observed. B) Long term potentiation (LTP) of patched CA1 pyramidal cells, peak amplitude of evoked EPSPs normalized to baseline recordings are plotted along the 35 minute timecourse in five minute bins for 6mo (right) and 14mo (left) animals. Inset with representative traces (Scale bar: 10ms, 20mV), 6mo 5XFAD *DLGAP2* overexpressing animals had diminished LTP compared to controls. C) Barplots showing dendritic spine density as number of thin, mushroom or stubby spines per 10μm of dendrite. Each dot represents a single cell, values from dendrites of that cell are averaged together with apical and basal denrites kept seperate. Plots are inset with rescaled y-axis to show effect sizes within mushroom and stubby spines. Apical thins spines are reduced in Ntg 6mo *DLGAP2* overexpressing mice compared to control AAV. D) Representative images and reconstructions from captured apical dendrites for each experimental group. Spine types are visualized by color (Yellow = Thin, Green = Mushroom, Red = Stubby, Blue = Filopodia). Scale bar = 5μm. E) Barplots showing dendritic spine density as number of thin, mushroom or stubby spines per 10μm of dendrite from cultured rat hippocampal neurons at 14 days in vitro. Each dot represents a single cell. Thins spines are reduced in cultures overexpressing Dlgap2 with no amyloid beta (n = 8-15 cells per group). p-values were derived from a t-test and corrected for multiple testing error using the Bonferroni method (*p<0.05, **p<0.01 ***p<0.001). Error bars are representad as standard error around the mean. n sizes for animals and cells are listed in Table 1.

To assess alterations in synaptic plasticity, we measured LTP across a 40-minute period. Two-way repeated measures ANOVA detected a significant main effect of AAV construct (F(1, 70) = 4.56, p < 0.05), as well as a significant age x AAV construct interaction (F(1, 70) = 7.94, p < 0.01). At 6mo, *DLGAP2* overexpression decreased potentiation compared to controls in 5XFAD animals; however, by 14mo, neither controls nor *DLGAP2* overexpressing mice exhibited LTP, regardless of AD status (Fig. 3B) possibly because there is an impairment floor effect that prevents measuring diminished LTP passed this point. To verify that *DLGAP2* overexpression was not acting presynaptically via overexpression in CA3 neurons, we measured Paired Pulse Ratio (PPR). We detected no significant effect of AAV construct, AD status or age on PPR (Fig. S7). These data suggest that overexpression of *DLGAP2* not only fails to strengthen synaptic throughput and LTP but also weakens these outcomes in the presence of AD pathology. There were no significant main effects of *DLGAP2* overexpression, age, or AD status on any of the other intrinsic measurements before or after TBS (Fig. S8), suggesting DLGAP2 has specific effects at synapse and not via general effects on neuron excitability.

### 3.5 DLGAP2 overexpression reduces thin spine density in young animals without AD pathology

To determine if the observed deficits in synaptic plasticity and subsequent cognitive outcomes from *DLGAP2* overexpression in AD mice can be explained by synaptic structure, we analyzed apical and basal dendritic spine morphologies of patched CA1 pyramidal cells. Spine density was largely comparable between apical and basal dendrites regardless of age, 5XFAD status and/or AAV construct. We observed significant main effects of AAV construct (F(1,293) = 13.84, p < 0.001) and age (F(1, 293) = 13.03, p < 0.001), as well as an AD status x age interaction (F(1, 293) = 5.71, p < 0.05), on dendritic spine density. Although 5XFAD animals showed no significant changes in spine density across any spine type at 6mo or 14mo, Ntg animals exhibited a decrease in thin spine density at 6mo of age compared to control injections (Fig. 3C-D). However, by 14 months, thin spine density was comparable to both 6 and14mo eGFP controls, confirming a role for *DLGAP2* in synaptic development and maturation of control mice, but is compensated throughout an animal’s lifespan. Interestingly, we see similar outcomes in cultured rat hippocampal cells; by DIV14, cells treated with Dlgap2 cDNA without the presence of amyloid beta have reduced thin spine densities (mean = 1.7, SD = 1.3) compared to GFP treated controls (mean = 2.7, SD = 1.1) (Fig. 3E, S9). In vitro results highlight the utility of cell culture experiments in studying postsynaptic proteins in a translationally relevant manner. All together however, these results indicate that the effects of DLGAP2 overexpression on behavior and synaptic plasticity in F1 hybrid AD mice are not explained by changes in spine morphology.

## 4. Discussion

This study builds upon our previous work and provides new insights into the role of *DLGAP2* overexpression in the hippocampus within the context of AD pathology. Overexpression of *DLGAP2* had unexpected effects on behavioral outcomes; where we expected overexpression to rescue cognitive deficits in an AD mouse model, we observed exacerbated cognitive decline. While fully understanding the mechanisms driving this effect will require further study, there are several putative mechanisms that may be responsible. Previous work has shown that overexpression of certain SHANK isoforms, a PSD protein that interacts with *DLGAP2*, leads to manic-like behavior, including increased locomotor activity and sleep disruption resultant from CNS excitatory/inhibitory imbalance [14]. While we did not measure sleep patterns in this study there are reports of sleep disruption in the 5XFAD mouse model [15, 16], which may explain the marked effects on locomotor activity and cognitive decline in 5XFAD animals that overexpress *DLGAP2*. However, if *DLGAP2* overexpression does affect sleep quality, the effect on cognition is unclear as this study was unable to determine whether it directly affects cognitive outcomes or if these outcomes result from disruption of sleep [17]. Similarly, overexpression of Homer1a, an interacting protein with *DLGAP2* and SHANK, in the dorsal hippocampus leads to impaired working memory in mice [18], suggesting that regulation of these PSD proteins has significant effects on cognitive outcomes. It has been proposed that overexpression of exogenous proteins may outcompete binding sites for necessary interacting proteins, leading to unexpected mutant phenotypes [19]. It is very possible that a similar phenomenon is occurring with *DLGAP2* in the PSD where there are a multitude of tightly regulated proteins interacting with each other. An influx of exogenous *DLGAP2* could very well outcompete other binding sites across the PSD leading to synaptic destabilization, especially in a vulnerable pre and post AD pathology state. Further validation is required to confirm this.

Given that AD pathology is known to cause dendritic spine loss [20, 21], it is interesting that we observed spine loss in 6mo Ntg animals overexpressing *DLGAP2* and not 5XFAD counterparts. There is clearly an unexpected interaction between AD status and *DLGAP2* overexpression in which pre-symptomatic AD pathology factors prevent the initial disruption of spine maturation caused by overexpression of *DLGAP2*. Since we see a loss of these spines only in apical and not basal dendrites, a possible explanation is that *DLGAP2* overexpression may hinder function of CA3 neurons resulting in a reduction of CA1 post synaptic activity and dendritic spine development in Ntg animals. Presymptomatic soluble Aβ has been shown to induce selective hyperexcitability in the hippocampus, which could overcome the deficits induced by *DLGAP2* overexpression [22–25], explaining why we see this effect in only Ntg animals. However, loss of thin spine density does not coincide with a significant change in spatial working memory or contextual fear related memory outcomes and is largely recovered by 14mo of age, limiting the biological significance of this result.

Hippocampal overexpression of *DLGAP2* had minimal effect on synaptic and intrinsic properties of CA1 pyramidal neurons. While we observed that 6mo 5XFAD animals overexpressing *DLGAP2* had diminished LTP compared to GFP injected controls, this effect was not present by 14mo and did not coincide with observed cognitive deficits. Based on the LTP results there is an apparent hyperexcitability phenotype in 6mo 5XFAD animals that is diminished in *DLGAP2* overexpressing animals and is comparable to age-matched Ntg controls. However, we did not observe a significant difference in synaptic throughput between *DLGAP2* overexpressing and control animals suggesting that the observed effects in LTP are not explained by synaptic hyperexcitability alone. It is important to note that expression was modest in CA1, with higher intensity observed in cells originating from CA3/Dentate/Subiculum and in presynaptic projections from recorded regions, as such, effects of *DLGAP2* overexpression on LTP in recorded CA1 neurons may act through presynaptic projections from CA3.

Our work has shown that viral mediated hippocampal overexpression of *DLGAP2* is not an effective means to ameliorate AD-related pathology. While the exact mechanism by which overexpression of *DLGAP2* negatively impacts AD-related cognition will require further investigation, we have shown that *DLGAP2* does play a role in AD-related cognitive outcomes.

## 6. Acknowledgments

We would like to acknowledge Nicholas Jewett for performing animal surgeries. Also, Niran Hadad and Surjeet Singh for their intellectual contributions to this work.

## 9. Consent Statement

No human subjects were part of this research.

## References

1. Ouellette, A.R., et al., Cross-Species Analyses Identify Dlgap2 as a Regulator of Age-Related Cognitive Decline and Alzheimer’s Dementia. Cell Reports, 2020. 32(9): p. 108091.

2. Shin, S.M., et al., GKAP orchestrates activity-dependent postsynaptic protein remodeling and homeostatic scaling. Nature Neuroscience, 2012. 15(12): p. 1655–1666.

3. Kajimoto, Y., et al., Synapse-Associated Protein 90/Postsynaptic Density-95-Associated Protein (SAPAP) is Expressed Differentially in Phencyclidine-Treated Rats and is Increased in the Nucleus Accumbens of Patients with Schizophrenia. Neuropsychopharmacology, 2003. 28(10): p. 1831–1839.

4. Pinto, D., et al., Functional impact of global rare copy number variation in autism spectrum disorders. Nature, 2010. 466(7304): p. 368–372.

5. Bienvenu, O.J., et al., Sapap3 and pathological grooming in humans: Results from the OCD collaborative genetics study. American Journal of Medical Genetics Part B: Neuropsychiatric Genetics, 2009. 150B(5): p. 710–720.

6. Neuner, S.M., et al., Harnessing Genetic Complexity to Enhance Translatability of Alzheimer’s Disease Mouse Models: A Path toward Precision Medicine. Neuron, 2019. 101(3): p. 399–411.e5.

7. Dunn, A.R., et al., A genetically-diverse mouse model reveals a complex gene-environment regulation of cognitive resilience and susceptibility to Alzheimer’s disease. bioRxiv, 2025: p. 2025.02.07.637137.

8. Erdfelder, E., F. Faul, and A. Buchner, *GPOWER: A general power analysis program.* Behavior Research Methods, Instruments & Computers, 1996. 28(1): p. 1–11.

9. Neuner, S.M., et al., TRPC3 channels critically regulate hippocampal excitability and contextual fear memory. Behavioural Brain Research, 2015. 281: p. 69–77.

10. Dunn, A.R., et al., Cell-Type-Specific Changes in Intrinsic Excitability in the Subiculum following Learning and Exposure to Novel Environmental Contexts. eNeuro, 2018. 5(6).

11. Neuner, S.M., et al., Harnessing Genetic Complexity to Enhance Translatability of Alzheimer’s Disease Mouse Models: A Path toward Precision Medicine. Neuron, 2019. 101(3): p. 399–411.e5.

12. Trott, J.M., et al., Conditional and unconditional components of aversively motivated freezing, flight and darting in mice. Elife, 2022. 11.

13. Gruene, T.M., et al., Sexually divergent expression of active and passive conditioned fear responses in rats. eLife, 2015. 4: p. e11352.

14. Han, K., et al., SHANK3 overexpression causes manic-like behaviour with unique pharmacogenetic properties. Nature, 2013. 503(7474): p. 72–7.

15. Sethi, M., et al., Increased fragmentation of sleep-wake cycles in the 5XFAD mouse model of Alzheimer’s disease. Neuroscience, 2015. 290: p. 80–9.

16. Drew, V.J., C. Wang, and T. Kim, Progressive sleep disturbance in various transgenic mouse models of Alzheimer’s disease. Front Aging Neurosci, 2023. 15: p. 1119810.

17. Lloret, M.A., et al., Is Sleep Disruption a Cause or Consequence of Alzheimer’s Disease? Reviewing Its Possible Role as a Biomarker. Int J Mol Sci, 2020. 21(3).

18. Celikel, T., et al., Select overexpression of homer1a in dorsal hippocampus impairs spatial working memory. Front Neurosci, 2007. 1(1): p. 97–110.

19. Prelich, G., Gene overexpression: uses, mechanisms, and interpretation. Genetics, 2012. 190(3): p. 841–54.

20. Boros, B.D., et al., Dendritic spines provide cognitive resilience against Alzheimer’s disease. Ann Neurol, 2017. 82(4): p. 602–614.

21. Mijalkov, M., et al., Dendritic spines are lost in clusters in Alzheimer’s disease. Scientific Reports, 2021. 11(1): p. 12350.

22. Busche, M.A., et al., Critical role of soluble amyloid-β for early hippocampal hyperactivity in a mouse model of Alzheimer’s disease. Proceedings of the National Academy of Sciences, 2012. 109(22): p. 8740–8745.

23. Ray, A., et al., Early hippocampal hyperexcitability and synaptic reorganization in mouse models of amyloidosis. iScience, 2024. 27(9): p. 110629.

24. Quiroz, Y.T., et al., Hippocampal hyperactivation in presymptomatic familial Alzheimer’s disease. Annals of Neurology, 2010. 68(6): p. 865–875.

25. Palop, J.J., et al., Aberrant Excitatory Neuronal Activity and Compensatory Remodeling of Inhibitory Hippocampal Circuits in Mouse Models of Alzheimer’s Disease. Neuron, 2007. 55(5): p. 697–711.

