## Supplemental Figures for "Evaluation of hippocampal *DLGAP2* overexpression on cognition, synaptic function, and dendritic spine structure in a translationally relevant AD mouse model"

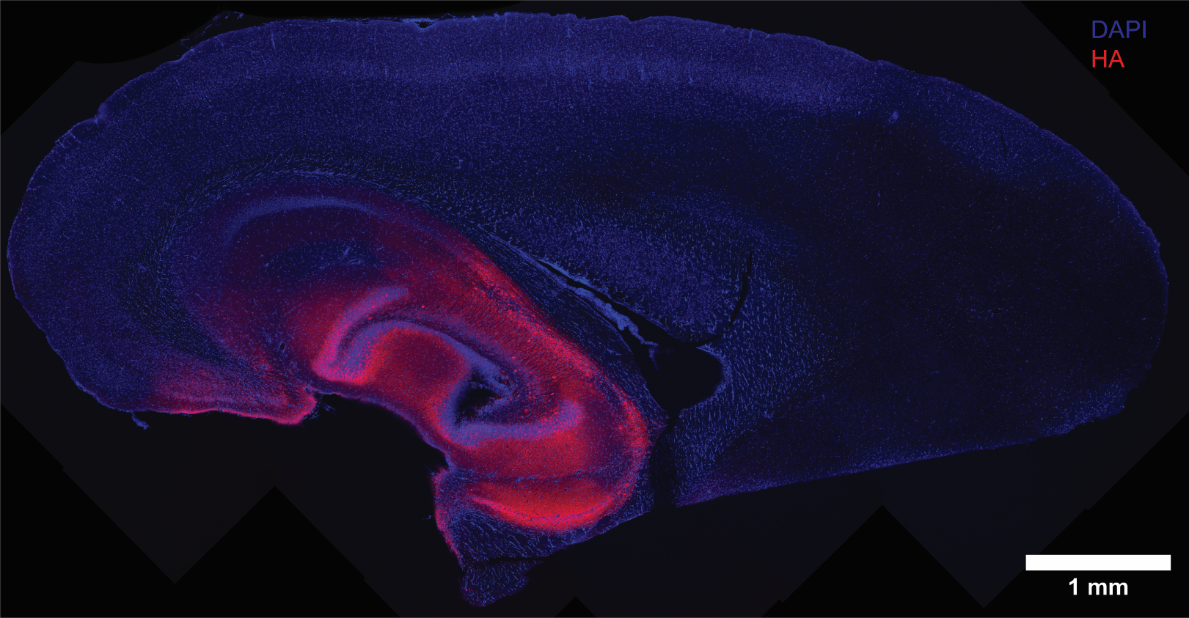

**Figure S1 | AAV spread is limited to hippocampus.** Representative image of a transverse brain section from a 14mo old animal injected with Dlgap2 overexpression AAV. Imaging of HA reporter shows that Dlgap2 overexpression is isolated to the hippocampus and entorhinal cortex. The cerebral cortex was not transfected. 45 Dlgap2 overexpressing samples were imaged.

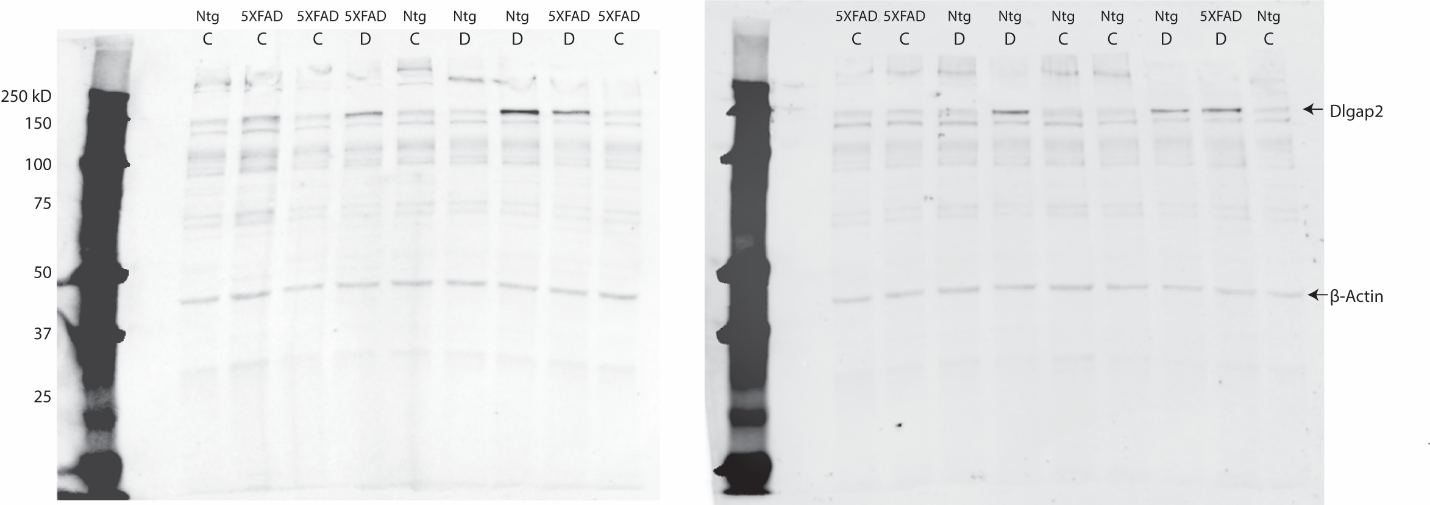

**Figure S2 | Western blot quantification of DLGAP2.** Raw images of western blots used to quantify Dlgap2 expression and loading control β-Actin in hippocampal samples overexpressing Dlgap2 (D) or control eGFP (C) from 14mo aged animals. Imaged at 700um wavelength with 10s exposure length. N= 3-5 per experimental group.

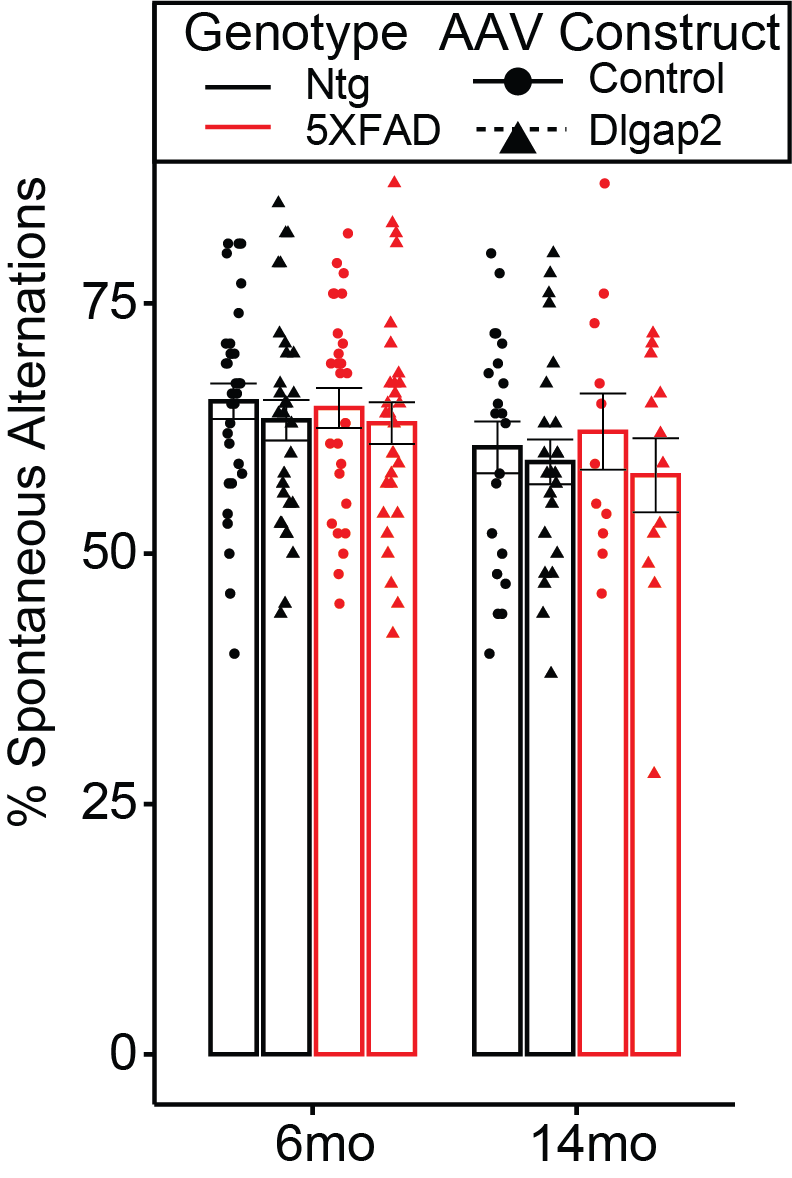

**Figure S3 | Y-maze working memory outcomes are unaffected by Dlgap2 overexpression.** Y-maze Spatial Working memory measured by % spontaneous alternations during 8 minutes of free exploration. There was no significant effect of Dlgap2 overexpression of 5XFAD status. Error bars are represented as standard error around the mean. n sizes for animals are listed in Table 1.

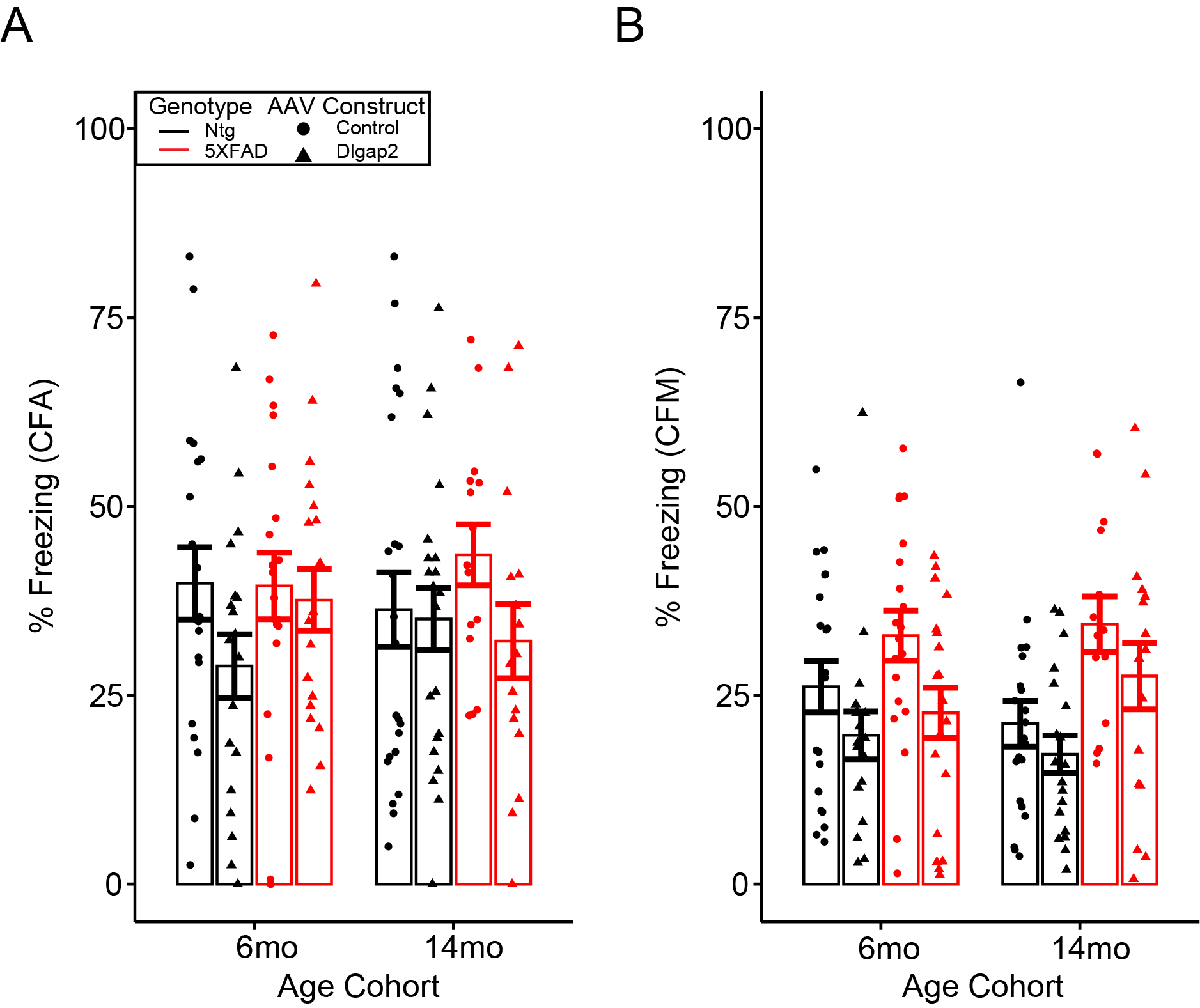

**Figure S4 | 6mo and 14mo cohorts have comparable cognitive outcomes at the initial 6mo timepoint.** A) Contextual fear acquisition measured (CFA) as % time spent freezing during the 40s post shock 4 (PS4) interval at 6mo of age. Data are shown for the 6mo and 14mo cohorts separately. B) 6mo contextual fear memory (CFM) recall measured by % time freezing during the testing session with no shock administration 24 hours after training. Error bars are represented as standard error around the mean.

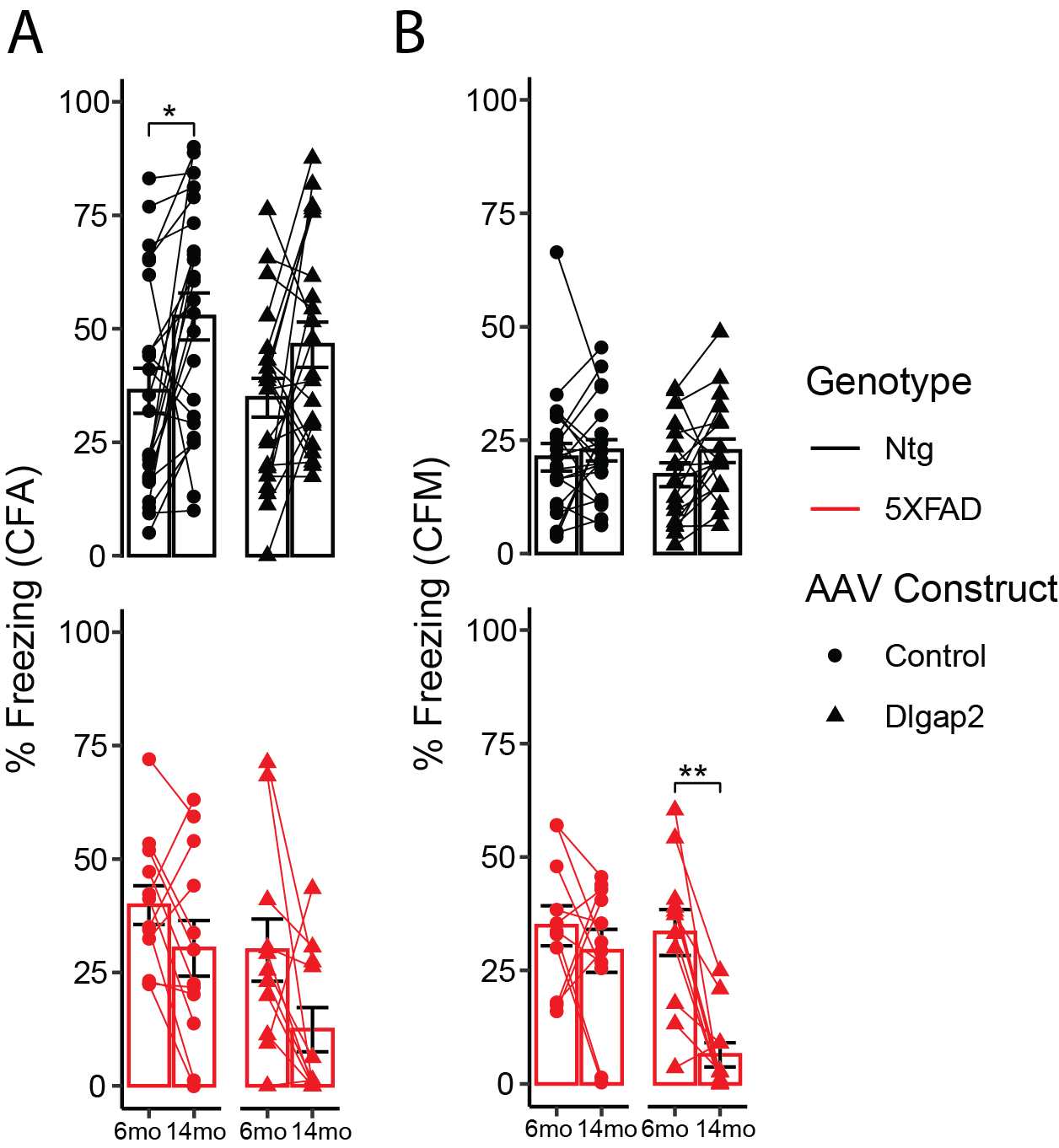

**Figure S5 | DLGAP2 overexpression and 5XFAD status coincide with age-related cognitive decline.** Cognitive outcomes for mice tested longitudinally at both 6 and 14mo for A) CFA and B) CFM. Age-related decline was strongest in 5XFAD animals overexpressing DLGAP2. Error bars are represented as standard error around the mean. p-values were derived from a t-test and corrected for multiple testing error using the Bonferroni method (*p<0.05, **p<0.01).

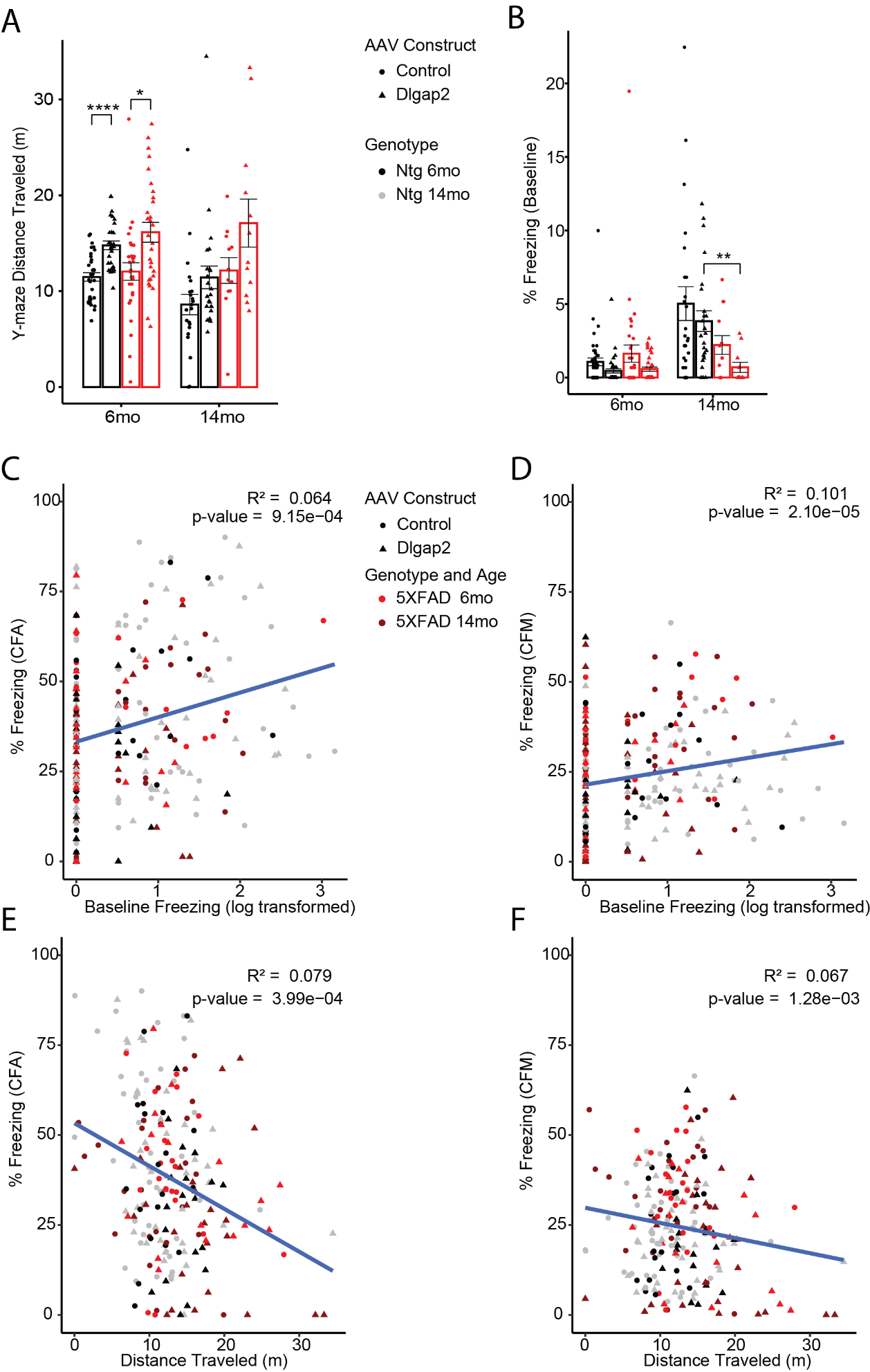

**Figure S6 | CFC outcomes correlate with locomotor measurements.** A) Total distance traveled in meters during the Y-maze spatial working memory task. B) Baseline freezing measured by % time freezing during training before shocks are administered. 5XFAD mice overexpressing Dlgap2 had higher levels of freezing than Ntg counterparts but not control AAV counterparts. Error bars are representad as standard error around the mean. p-values were derived from a t-test and corrected for multiple testing error using the Bonferroni method (*p<0.05, **p<0.01 ***p<0.001). C) Correlation dot plot with log transformed baseline % freezing on the x-axis and contextual fear acquisition (PS4 % Freezing) on the y-axis. D) Correlation dot plot with log transformed baseline % freezing on the x-axis and contextual fear memory (Total % freezing on test day) on the y-axis. E) Correlation dot plot with distance traveled during Y-maze exploration on the x-axis and contextual fear acquisition (PS4 % Freezing) on the y-axis. F) Correlation dot plot with distance traveled during Y-maze exploration and contextual fear memory (Total % freezing on test day) on the y-axis. n sizes for animals are listed in Table 1.

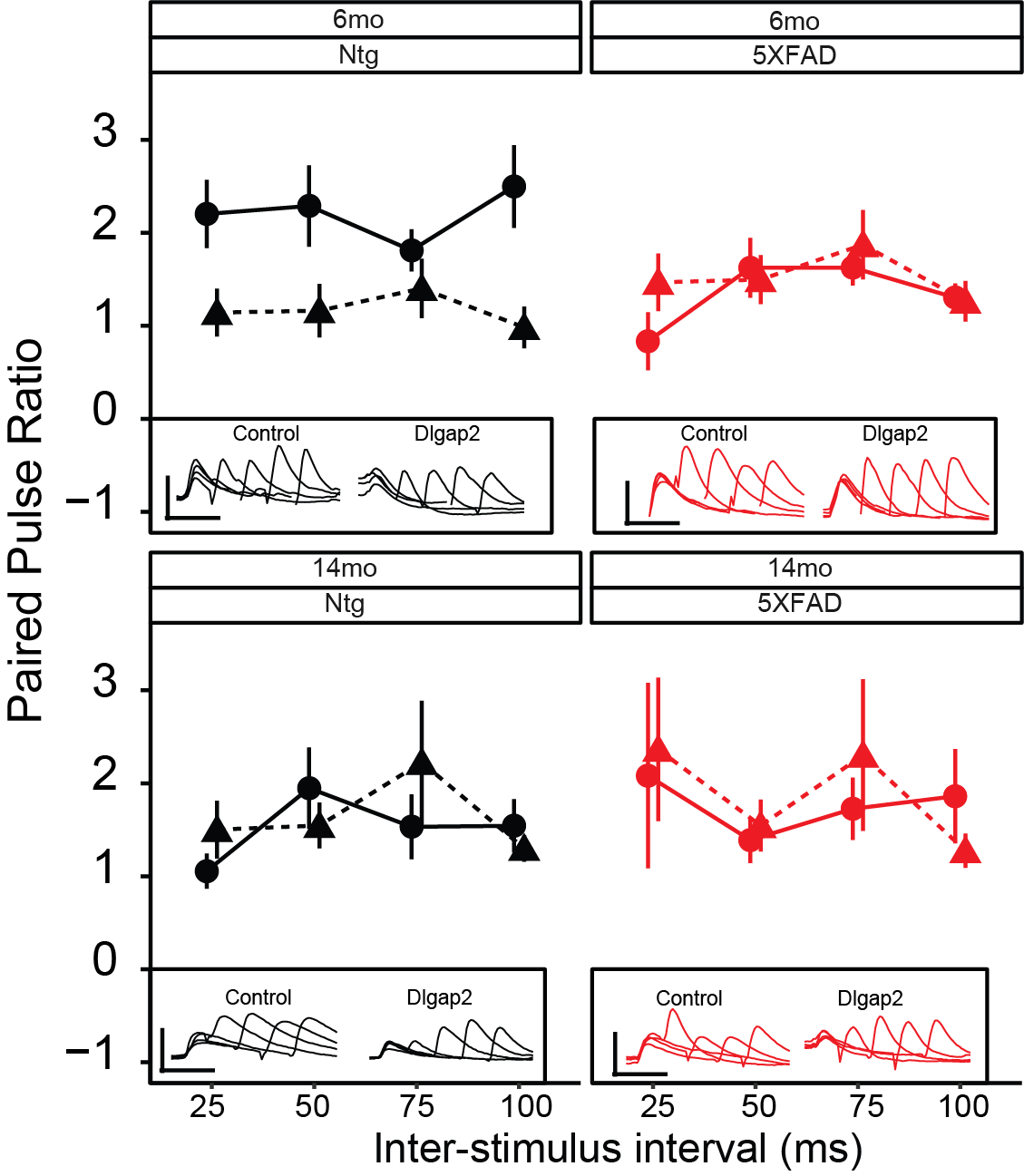

**Figure S7 | DLGAP2 overexpression does not affect paired pulse ratio.** Paired Pulse Ratio measured across four pulse intervals. Inset with representative traces (Scale bar: 50ms, 10mV). p-values were derived from a t-test and corrected for multiple testing error using the Bonferroni method (*p<0.05, **p<0.01 ***p<0.001). Error bars are representad as standard error around the mean. n sizes for animals and cells are listed in Table 1.

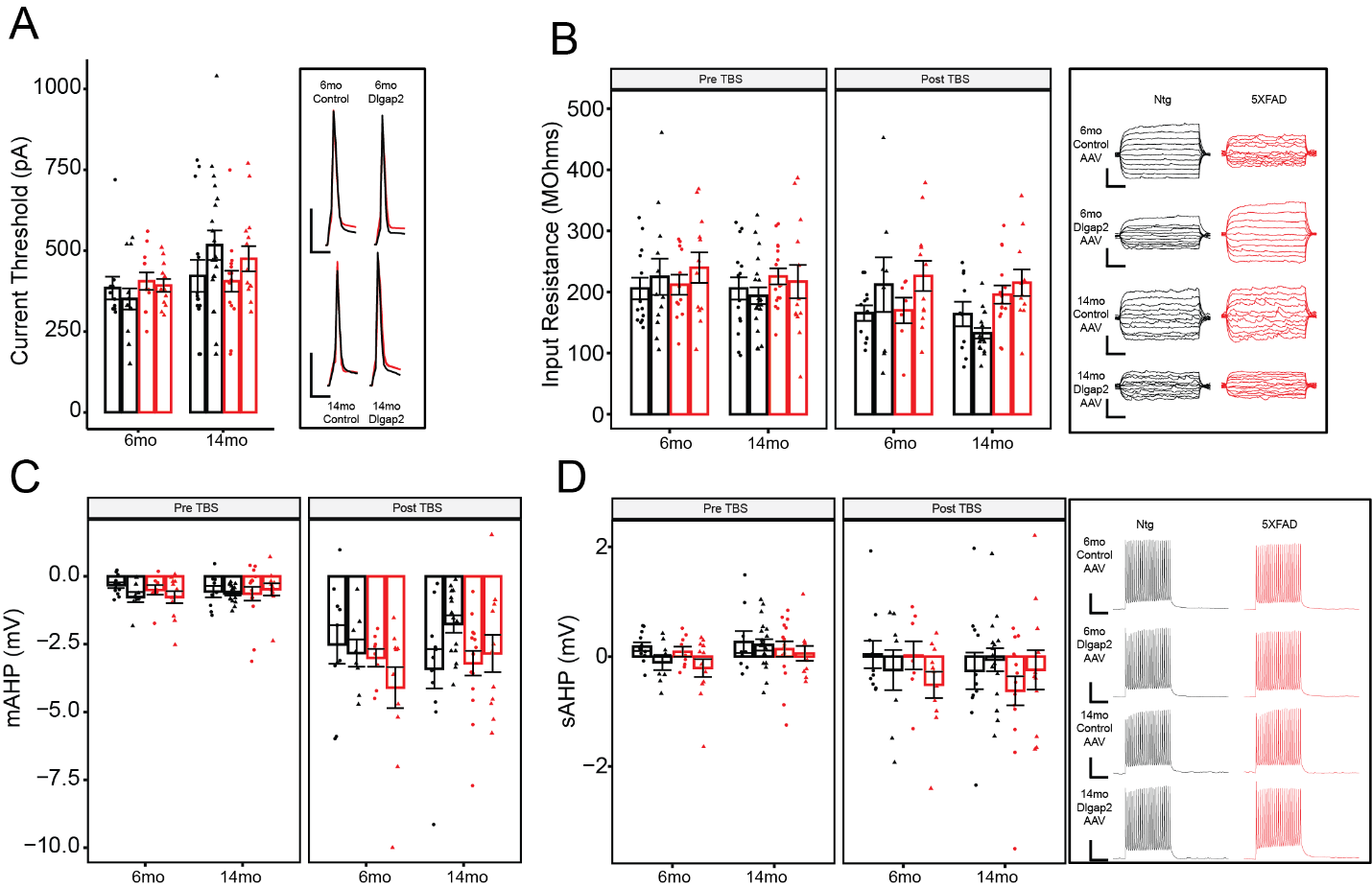

**Figure S8 | DLGAP2 overexpression does not affect intrinsic firing properties.** A) Action potential current threshold with representative traces. (Scale bar: 10ms, 40mV). B) Input Resistance measured as change in membrane potential (mV) in response to varying stimulating current ranging from -50pA to 50pA, along with representative traces (Scale bar: 250ms, 10mV). C) Medium after-hyperpolarization (mAHP). D) Slow after-hyperpolarization (sAHP) (Scale bar: 200ms,25mv). No main effect of DLGAP2 overexpression was observed in C-G. Error bars are displayed as standard error around the mean. p-values were derived from a t-test and corrected for multiple testing error using the Bonferroni method (*p<0.05, **p<0.01 ***p<0.001). n sizes for animals and cells are listed in Table 1.

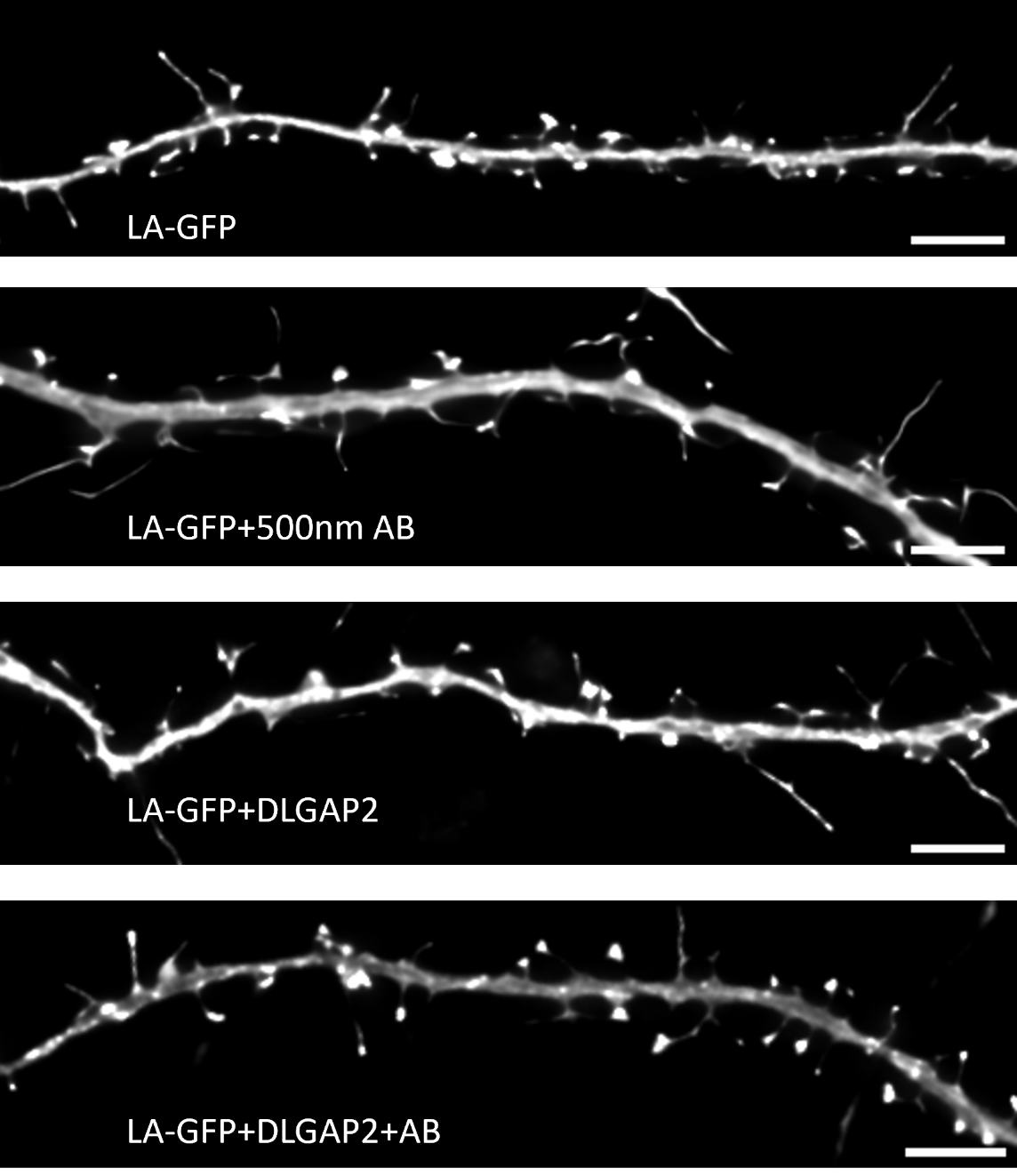

**Figure S9 | In vitro dendritic spine representative images.** Representative images from 14 DIV cultured rat hippocampal neurons for each experimental group. Scale bar = 5μm.
